# Nuclear receptor signaling via NHR-49/MDT-15 regulates stress resilience and proteostasis in response to reproductive and metabolic cues

**DOI:** 10.1101/2023.04.25.537803

**Authors:** Ambre J. Sala, Rogan A. Grant, Ghania Imran, Claire Morton, Renee M. Brielmann, Laura C. Bott, Jennifer Watts, Richard I. Morimoto

## Abstract

The ability to sense and respond to proteotoxic insults declines with age, leaving cells vulnerable to chronic and acute stressors. Reproductive cues modulate this decline in cellular proteostasis to influence organismal stress resilience in *C. elegans*. We previously uncovered a pathway that links the integrity of developing embryos to somatic health in reproductive adults. Here, we show that the nuclear receptor NHR-49, a functional homolog of mammalian peroxisome proliferator-activated receptor alpha (PPARα), regulates stress resilience and proteostasis downstream of embryo integrity and other pathways that influence lipid homeostasis, and upstream of HSF-1. Disruption of the vitelline layer of the embryo envelope, which activates a proteostasis-enhancing inter-tissue pathway in somatic tissues, also triggers changes in lipid catabolism gene expression that are accompanied by an increase in fat stores. NHR-49 together with its co-activator MDT-15 contributes to this remodeling of lipid metabolism and is also important for the elevated stress resilience mediated by inhibition of the embryonic vitelline layer as well as by other pathways known to change lipid homeostasis, including reduced insulin-like signaling and fasting. Further, we show that increased NHR-49 activity is sufficient to suppress polyglutamine aggregation and improve stress resilience in an HSF-1-dependent manner. Together, our results establish NHR-49 as a key regulator that links lipid homeostasis and cellular resilience to proteotoxic stress.

## Introduction

Maintaining a properly folded and functional proteome is essential for cellular and organismal health (Labbadia and Morimoto 2015a; Sala et al. 2017). The capacity of cells to prevent the formation and accumulation of misfolded and aggregated protein species, known as protein homeostasis (proteostasis), declines over time during aging which is considered a major driver of age-dependent cellular dysfunction (Hipp et al. 2019). Abnormal protein aggregates are observed in many age-related disorders, including dementia, diabetes, and muscular dystrophy (Balch et al. 2008; Wilson et al. 2023). To counteract protein misfolding and aggregation, cells have evolved a sophisticated network of cellular machineries for protein synthesis, folding, and clearance, consisting mostly of molecular chaperones and protein degradation pathways (Labbadia and Morimoto 2015a; Sala et al. 2017). Proteostasis is constantly challenged by physiological and environmental stresses, and cells are also equipped with adaptive mechanisms that are activated during proteotoxic stress. Loss of proteostasis in the cytosol activates the heat shock response (HSR) which triggers the rapid and transient induction of heat shock proteins (HSPs), or chaperones, and is mainly regulated by the evolutionarily conserved heat shock transcription factor 1 (HSF1) (Gomez-Pastor et al. 2018).

In addition to surveillance mechanisms at the cellular level, proteostasis regulation at the systemic level involves responses to physiological and metabolic cues that determine organismal health during aging (Hansen et al. 2013; Li et al. 2017; Sala et al. 2017). Rewiring of metabolism towards survival is generally associated with improved proteostasis, for example in conditions where insulin signaling is reduced (Hsu et al. 2003; Morley and Morimoto 2004; Hansen et al. 2013). Mutations or treatments that impair reproduction have also been shown to promote somatic resilience in several model systems (Antebi 2013). Removal of germ cells in *C. elegans* and *Drosophila* increases lifespan and suppresses the age-dependent aggregation of disease model proteins in somatic tissues (Hsin and Kenyon 1999; Flatt et al. 2008; Shemesh et al. 2013). In wild type *C. elegans*, signals from germline stem cells (GSC) initiate the collapse of proteostasis that occurs at reproductive maturity via the generalized repression of the HSR, possibly to prioritize reproduction at the expense of somatic health (Shemesh et al. 2013; Labbadia and Morimoto 2015b). However, somatic resilience and reproduction can be uncoupled, and specific molecular pathways rather than mere energetic trade-offs govern the regulation of proteostasis mechanisms by the reproductive system (Antebi 2013; Labbadia and Morimoto 2015b; Sala et al. 2020).

We recently identified a pathway in which signals emanating from fertilized eggs modulate proteostasis in the somatic tissues of reproductive mothers to promote maternal health when reproduction is compromised (Sala et al. 2020). We demonstrated that perturbation of the fertilized egg in the uterus, induced by inhibition of *cbd-1* or other genes encoding vitelline layer components, improves stress resilience and restores the HSR in maternal somatic tissues. This pathway involves a DAF-16/FOXO response specifically in the egg laying apparatus (vulva), which may be involved in monitoring egg quality, and relies on HSF-1, the master regulator of the HSR, in somatic tissues (Sala et al. 2020). The beneficial effects observed in response to alteration of the embryonic vitelline layer may have evolved to maximize maternal survival until conditions for optimal reproduction resume (Sala and Morimoto 2021). This relationship between the health of the fertilized egg and the health of maternal somatic tissues provides a paradigm for how systemic regulations influence organismal robustness and resilience.

Here, we report that the nuclear receptor NHR-49, a functional homolog of mammalian peroxisome proliferator-activated receptor alpha (PPARα), together with its coactivator the mediator subunit MDT-15, is an important regulator of lipid metabolism and stress resilience in response to alteration of the embryonic vitelline layer. Further, we find that NHR-49 plays a role in multiple paradigms of stress resilience that also alter lipid homeostasis, including reduced insulin signaling and fasting, and that increased NHR-49 activity is sufficient to improve stress resilience, the HSR and proteostasis, in an HSF-1-dependent manner. Together, our results demonstrate that NHR-49/PPARα has a pervasive role as a regulator that connects lipid homeostasis and cellular resilience to proteotoxic stress.

## Results

### Reduced function of the vitelline gene *cbd-1* has systemic effects on gene expression and upregulates lipid metabolism genes

We previously uncovered that alteration of the vitelline layer of fertilized embryos in the uterus initiates a transcellular pathway that results in improved maternal proteostasis and stress resilience (Sala et al. 2020). The *cbd-1(ok2913)* hypomorph mutation recapitulates the beneficial effects of vitelline inhibition on organismal stress resilience as shown by increased survival following a lethal heat stress at 35°C (Fig. 1A), while it also maintains the ability to produce a limited amount of viable progeny (Fig. 1B) (Johnston et al. 2010; Sala et al. 2020). To gain a better understanding of the systemic effects of vitelline inhibition, we asked how the *cbd-1(ok2913)* mutation affects gene expression in basal conditions and following a heat stress using RNA sequencing (Fig. S1A).

**Figure 1.**
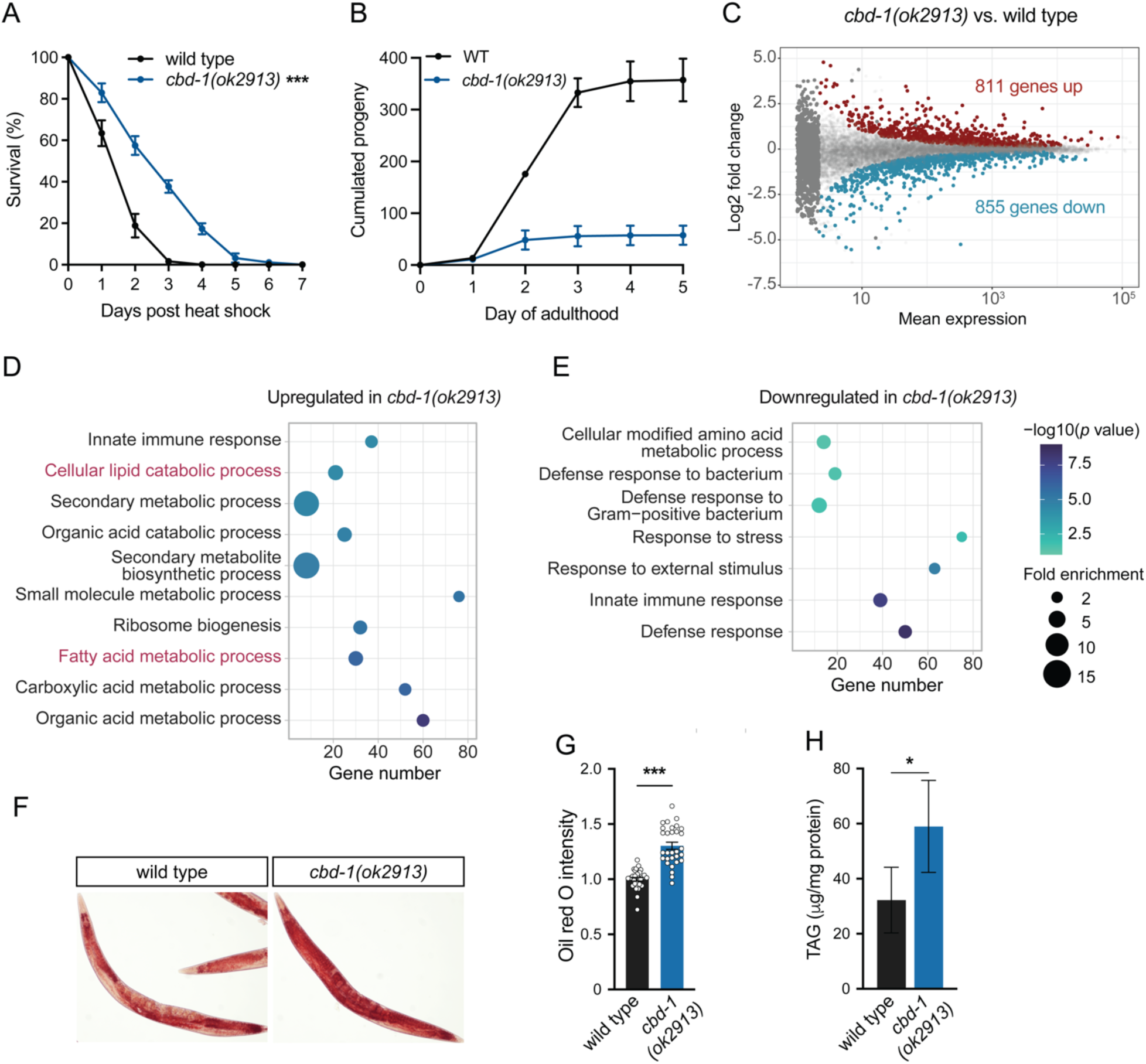
Lipid metabolism is altered in *cbd-1(ok2913)* (A) Heat stress survival of wild type or *cbd-1(ok2913)* animals following a 35 °C heat shock for 4 hours on day 2 of adulthood and allowed to recover at 20 °C (*n* = 3). (B) Cumulated number of progeny produced by wild type or *cbd-1(ok2913)* animals at 20°C through day 5 of adulthood. Wild type *n* = 10, *cbd-1(ok2913) n* = 12. (C) Differentially expressed genes in *cbd-1(ok2913)* versus wild type animals as determined by RNA sequencing. Red, upregulated. Blue, downregulated. (D-E) Gene Ontology (GO) enrichment analysis of genes differentially upregulated (D) or downregulated (E) in *cbd-1(ok2913)*. The top 10 GO terms are shown. (F) Representative micrographs of oil red O-stained wild type and *cbd-1(ok2913)* animals. (G) Quantification of excess red signal in oil red O-stained micrographs. *n* = 30. (H) Quantification of triglycerides (TAG) in whole extracts of wild type and *cbd-1(ok2913)* animals. Error bars represent SEM. Statistical significance based on two-way ANOVA (A) or unpaired *t*-test (G, H). (*) *P* < 0.05; (***) *P* < 0.001.

Differential gene expression analysis identified 811 upregulated and 855 downregulated genes (*q* < 0.05, Wald test) in *cbd-1(ok2913)* compared to wild type animals (Fig. 1C), indicating a broad remodeling of transcription. Tissue enrichment analysis revealed that the differentially expressed genes were enriched for transcripts preferentially expressed in the intestine and other somatic tissues, confirming that although the *cbd-1* gene is restricted to the reproductive system (Johnston et al. 2010; Gonzalez et al. 2018; Sala et al. 2020), modulating its activity has widespread organismal consequences (Fig. S1B). Gene ontology (GO) analysis indicated an enrichment of genes related to innate immunity, ribosome biogenesis, and metabolism, in particular lipid metabolism, in the set of differentially upregulated genes in *cbd-1(ok2913)* (Fig. 1D). Downregulated genes were characterized by GO terms mostly related to innate immunity and defense responses (Fig. 1E). Genes differentially induced in *cbd-1(ok2913)* specifically during heat shock were similarly characterized by GO terms related to lipid metabolism and innate immunity (Fig. S1C). Importantly, the canonical heat shock gene *hsp-70* was among the genes more highly induced by heat stress in the *cbd-1(ok2913)* mutant (Fig. S1D), in agreement with our previous findings (Sala et al. 2020). Enrichment analysis of the phenotypes associated with the genes differentially expressed in the *cbd-1* mutant revealed that in addition to stress sensitivity, these genes are involved in regulating fat content (Fig. S1E), further suggesting that lipid metabolism may be affected in response to compromised eggs.

### Inhibition of the embryonic vitelline layer increases fat storage and triglyceride levels

Lipid metabolism has been shown to be altered in animals lacking GSCs and in conditions of reduced insulin signaling that are also associated with elevated proteostasis and stress resilience (Wang et al. 2008; O’Rourke et al. 2009). In both cases, the mutants accumulate fat stores compared to wild type animals (O’Rourke et al. 2009). To assess whether fat storage is altered in response to inhibition of the embryonic vitellin layer, we used the lysochrome diazo dye oil red O (ORO) to stain neutral triglycerides and lipids. We consistently observed an accumulation of fat stores in adult *cbd-1* mutant compared to wild type animals (Fig. 1F, G). We also measured triglyceride levels in whole animal lysates and found an increase of ~1.8 fold between *cbd-1* mutant and wild type animals (Fig. 1H). Elevated fat storage is associated with a remodeling of fatty acid (FA) composition in animals lacking GSCs (Ratnappan et al. 2014; Amrit et al. 2016) and mutants with reduced insulin signaling (Shmookler Reis et al. 2011; Shi et al. 2013). To determine whether excess fat storage in the *cbd-1(ok2913)* mutant is also accompanied by changes in FA profile, we examined FA composition in whole animals using gas chromatography-mass spectrometry (GC-MS). We found that *cbd-1(ok2913)* and wild type animals had a similar FA composition profile at different stages of adulthood (Fig S2A, B). Together, this data indicates that changes in the expression of lipid metabolism genes in response to inhibition of the embryonic vitelline layer are accompanied by an increase in fat storage and total triglyceride levels, without remodeling of FA composition.

### Peroxisomal FA β-oxidation limits fat accumulation and contributes to stress resilience

To further understand the changes in lipid metabolism that occur when the embryonic vitelline layer is altered, we performed a KEGG pathway enrichment analysis on the gene set upregulated in *cbd-1(ok2913)* and found that FA degradation, peroxisomes, and lysosomes, were among the most enriched categories (Fig. 2A). This suggests that the mobilization of FA from triglycerides could be higher in the fat store accumulating *cbd-1(ok2913)* animals. Two lysosomal lipases, which break down triglycerides into free FA, as well as many of the genes encoding the key enzymes of the peroxisomal FA β-oxidation pathway involved in the breakdown of FA molecules, are consistently upregulated in *cbd-1(ok2913)* compared to wild type animals (Fig. 2B). While several steps of the FA β-oxidation pathway are controlled by redundant enzymes, *daf-22* encodes the unique peroxisomal Thiolase in *C. elegans*, the last enzyme in the β-oxidation pathway. We used RNA interference (RNAi) to knockdown the expression of *daf-22* and examine the effects of peroxisomal FA β-oxidation inhibition in the *cbd-1* mutant. We found that *daf-22* RNAi leads to an increase in ORO staining in *cbd-1(ok2913)* compared to the control RNAi condition (Fig. 2C, D), suggesting that upregulation of peroxisomal FA β-oxidation may serve to limit further accumulation of fat stores.

**Figure 2.**
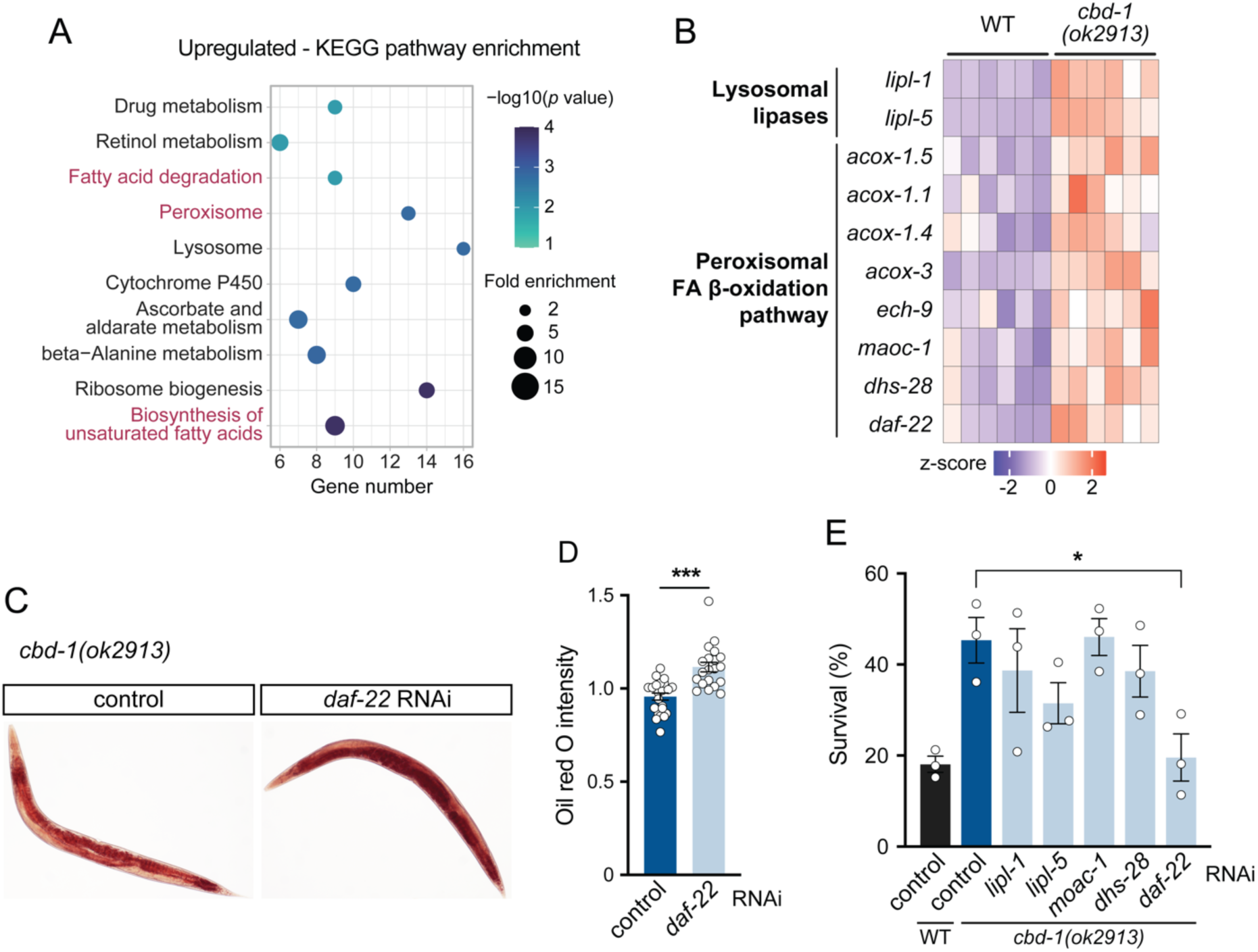
Peroxisomal FA β-oxidation limits fat accumulation and contributes to stress resilience. (A) KEGG pathway analysis of genes differentially upregulated in *cbd-1(ok2913)*. The top 10 terms are shown. (B) Relative expression of selected lipid catabolism genes in wild type (WT) and *cbd-1(ok2913)* animals. (C) Representative micrographs of oil red O-stained *cbd-1(ok2913)* animals grown on either control or *daf-22* RNAi. (D) Quantification of excess red signal in oil red O-stained micrographs. Control, *n* = 23; *daf-22* RNAi, *n* = 19. (E) Heat stress survival of wild type (WT) or *cbd-1(ok2913)* animals grown on the indicated RNAi and scored after 72 hours of recovery at 20 °C (*n* = 3). Error bars represent SEM. Statistical significance based on unpaired *t*-test (D) or one-way ANOVA followed by Dunnett correction (E). (*) *P* < 0.05; (***) *P* < 0.001.

We then asked whether these changes in lipid catabolism are important for the beneficial effects of inhibition of the embryonic vitelline layer on stress resilience. We used RNAi to knockdown the expression of lysosomal lipase and peroxisomal FA β-oxidation genes and analyzed the effects on stress resilience in *cbd-1(ok2913)*. We found that among the lipid catabolism genes tested, only *daf-22* RNAi suppressed the elevated stress resilience of *cbd-1(ok2913)* animals (Fig. 2E). While the negative results may be explained by functional redundancy, our finding with *daf-22* suggests that increased lipid catabolism via the peroxisomal FA β-oxidation pathway contributes to the elevated stress resilience induced by alteration of the embryonic vitelline layer.

### Vitelline disruption alters lipid metabolism genes through NHR-49 and its co-activator MDT-15

To identify mediators of the transcriptional changes observed in *cbd-1(ok2913)* mutants, we examined known regulators of lipid catabolism. Many of the genes differentially regulated in *cbd-1(ok2913)* are also responsive to the loss of the NHR-49 nuclear receptor, the functional homolog of human PPARα, that regulates FA β-oxidation during development and in conditions of food deprivation (Van Gilst et al. 2005a; Van Gilst et al. 2005b). Indeed, we observed that among the transcripts upregulated in *cbd-1(ok2913)*, 62 are also downregulated in the loss-of-function mutant *nhr-49(nr2041)* (*p* = 2.6 x 10^-22^, Fisher’s exact test) (Watterson et al. 2022) (Fig. 3A). This overlapping gene set primarily contained genes involved in FA metabolic processes, as shown by GO enrichment analysis (Fig. 3B), including genes encoding key enzymes of the peroxisomal FA β-oxidation pathway (*maoc-1*, *dhs-28*, *daf-22*). We thus tested whether NHR-49 as well as its known interactor and co-activator, the mediator subunit MDT-15 (Taubert et al. 2006; Goh et al. 2018), regulate the expression of lipid metabolism genes in *cbd-1* mutants. We selected two lipid metabolism genes that were among the top upregulated genes in *cbd-1(ok2913)*, and also downregulated in *nhr-49(nr2041)*, namely *far-3*, which encodes a fatty acid- and retinol-binding protein, and the cytochrome P450 encoding gene *cyp-29A2*. Real-time quantitative PCR (qPCR) analysis confirmed that mRNA levels of *far-3* and *cyp-29A2* were upregulated >10- and 3-fold, respectively, in *cbd-1(ok2913)* compared to wild type animals (Fig. 3C, D). Knockdown of *nhr-49* or *mdt-15* with RNAi blocked the induction of *far-3* (Fig. 3C) and *cyp-29A2* (Fig. 3D), indicating that the upregulation of these genes in *cbd-1(ok2913)* is strictly dependent on both NHR-49 and MDT-15. We then asked whether NHR-49 signaling also mediates the changes in fat stores and found that elevated fat storage in *cbd-1(ok2913)* was not affected by *nhr-49* loss-of-function mutation (Fig. 3E), indicating that NHR-49 activity is dispensable for this phenotype. Together, these results indicate that the NHR-49/MDT-15 nuclear receptor complex acts downstream of the increase in fat storage to mediate changes in lipid metabolism gene expression in response to inhibition of the embryonic vitelline layer.

**Figure 3.**
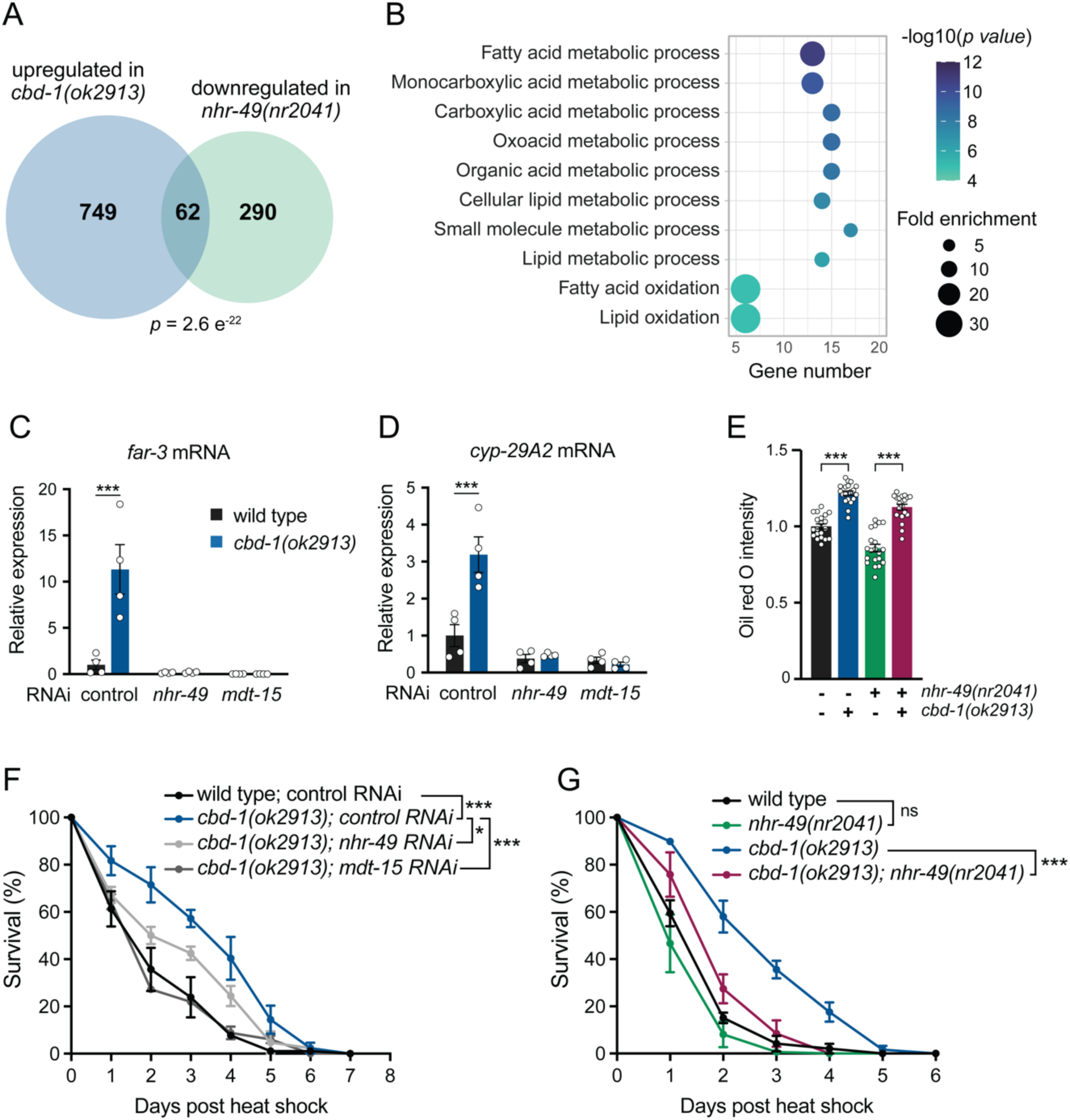
NHR-49 and its coactivator MDT-15 regulate lipid metabolism and stress resilience in *cbd-1(ok2913)* (A) Venn diagram showing the overlap between the set of genes upregulated in *cbd-1(ok2913)* and genes downregulated in *nhr-49(nr2041)* as determined in Watterson *et al* (2022). (B) GO enrichment analysis of the overlapping gene set shown in A. The top 10 GO terms are shown (C) Expression of *far-3* relative to *rpb-2* in day 2 adults normalized to wild type animals grown on control RNAi (*n* = 4). (D) Expression of *cyp-29A2* relative to *rpb-2* in day 2 adults normalized to wild type animals grown on control RNAi (*n* = 4). (E) Quantification of excess red signal in oil red O-stained micrographs for the indicated crosses. The ‘-‘ symbol denotes wild type allele for the corresponding locus (*n* = 20). (F) Heat stress survival of wild type or *cbd-1(ok2913)* animals grown on the indicated RNAi (*n* = 3). (G) Heat stress survival of wild type animals and indicated genotypes (*n* = 3). Error bars represent SEM. Statistical significance based on two-way ANOVA followed by Sidak correction (C, D), one-way ANOVA followed by Tukey correction (E), or two-way ANOVA (F, G). (*) *P* < 0.05; (***) *P* < 0.001, (ns [nonsignificant]) *P* > 0.05.

### NHR-49 and MDT-15 are required for the enhanced stress resilience in response to vitelline disruption

In addition to its role in regulating lipid metabolism, NHR-49 has previously been shown to also mediate resistance to oxidative stress, hypoxia, and pathogens (Goh et al. 2018; Wani et al. 2021; Doering et al. 2022). We then asked whether NHR-49/MDT-15 nuclear receptor signaling also plays a role in the elevated heat stress resilience induced by inhibition of the vitelline layer. We found that RNAi knockdown of *mdt-15* completely suppressed and *nhr-49* significantly reduced the prolonged survival of *cbd-1(ok2913)* following a lethal heat stress (Fig. 3F). The partial suppression of *nhr-49* was likely due incomplete penetrance of the RNAi since the loss-of-function allele *nhr-49(nr2041)* nearly completely abrogated the beneficial effects of *cbd-1(ok2913)* on heat stress survival (Fig. 3F). However, *nhr-49* loss-of-function had no significant effect on heat stress survival in the wild type background (Fig. 3G), which is in contrast to what has been reported for resistance to oxidative stress and hypoxia (Goh et al. 2018; Doering et al. 2022). Therefore, regulation by the NHR-49/MDT-15 nuclear receptor complex is required for the beneficial organismal effects in response to reproductive cues emanating from compromised embryos.

### NHR-49 is important in multiple paradigms of stress resilience

Considering the function of NHR-49 as a lipid-sensing factor that regulates gene expression, we hypothesized that it may have a general role in promoting stress resilience in diverse contexts where lipid homeostasis is altered. Therefore, we asked whether *nhr-49* loss-of-function influences heat stress survival in other paradigms of stress resilience known to also alter lipid homeostasis. Reduced activity of the insulin signaling pathway enhances resilience to multiple stresses and extends lifespan (Kenyon et al. 1993; Tatar et al. 2003). In *C. elegans*, these effects are mediated by the DAF-16/FOXO transcription factor and also require HSF-1, downstream of the insulin receptor DAF-2 (Hsu et al. 2003; Morley and Morimoto 2004). Reduced insulin signaling also results in accumulation of fat stores as measured by ORO staining and triglycerides quantification (O’Rourke et al. 2009). As expected, reduced insulin signaling in *daf-2(e1370)* hypomorphic mutants strongly enhanced survival following a lethal heat stress (Fig. 4A). Combining the *nhr-49(nr2041*) loss-of-function allele with *daf-2(e1370)* resulted in a substantial suppression of heat stress survival (Fig. 4A), indicating that NHR-49 contributes to stress resilience during reduced insulin signaling.

**Figure 4.**
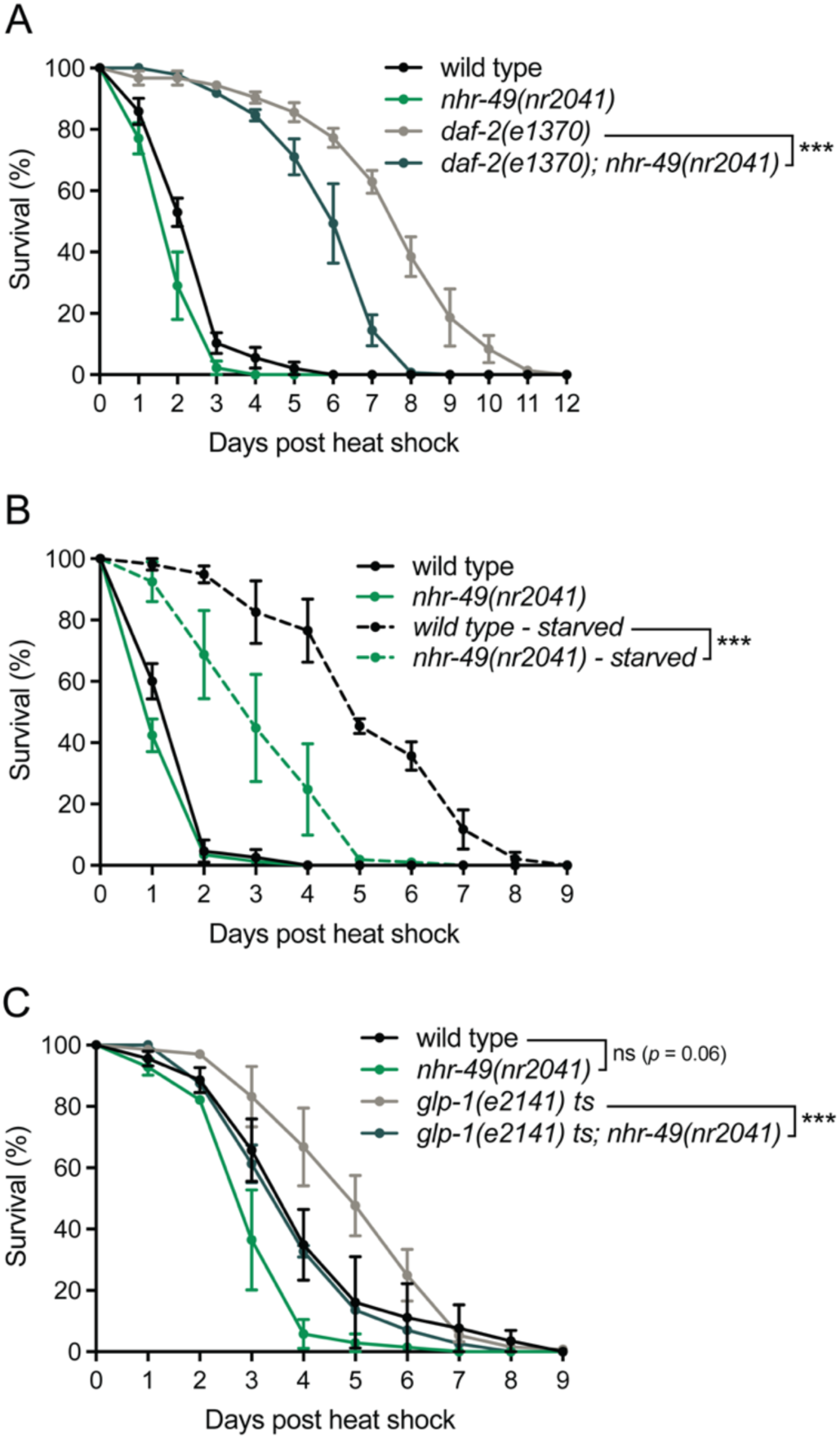
NHR-49 is important in multiple paradigms of stress resilience. (A) Heat stress survival of wild type and *daf-2(e1370)* mutants crossed into *nhr-49(nr2041)* (*n* = 3). (B) Heat stress survival of wild type and *nhr-49(nr2041)* grown on OP50 bacteria (fed) or deprived of food source for 16 hours prior to the heat stress (starved) (*n* = 3). (C) Heat stress survival of wild type and *glp-1(e2141)* temperature sensitive mutants crossed into *nhr-49(nr2041)*. Animals were grown at 25°C prior to heat stress and transferred back to 25°C (*n* = 3). Error bars represent SEM. Statistical significance based on two-way ANOVA. (***) *P* < 0.001, (ns [nonsignificant]) *P* > 0.05.

Reduced food intake ameliorates lifespan, proteostasis, and stress resilience in many organisms (Fontana and Partridge 2015). NHR-49 and its mammalian counterpart PPARα are important regulators of lipid metabolism genes in response to fasting to promote the utilization of fat stores (Leone et al. 1999; Van Gilst et al. 2005b). Therefore, we asked whether NHR-49 also contributes to the beneficial effects of fasting on stress resilience. As expected, wild type animals subjected to fasting for 16 hours prior to exposure to a lethal heat stress exhibited extended survival compared to the fed control (Fig. 4B). The effect of fasting on heat stress survival was reduced by about 50% in *nhr-49(nr2041*) animals (Fig. 4B), indicating that NHR-49 also regulates stress resilience in response to reduced food intake.

Loss of GSCs extends lifespan in an NHR-49-dependent manner and is known to result in increased lipid catabolism and accumulation of fat stores (O’Rourke et al. 2009; Ratnappan et al. 2014). As expected, *glp-1(e2141)* temperature sensitive mutants raised at the non-permissive temperature of 25°C exhibited elevated survival to a lethal heat stress as compared to wild type animals (Fig. 4C). Similar to what we observed for insulin signaling and fasting, *nhr-49* loss-of-function reduced the elevated stress resilience of *glp-1(e2141)* (Fig. 4C). In these conditions, we also observed a more pronounced but non-significant (p = 0.06) deleterious effect of *nhr-49* loss-of-function in the wild type background (Fig. 4C) compared to experiments where the animals were grown at the standard cultivation temperature of 20°C (Fig. 3G, Fig.4A, B). Together, these results indicate that NHR-49 has a pervasive role in stress resilience in conditions in which lipid metabolism is also altered.

### Increased NHR-49 activity enhances organismal stress resilience and proteostasis

Since NHR-49 mediates stress resilience in several contexts, we asked whether increasing the activity of this nuclear receptor was sufficient to enhance stress resilience. To test this, we first used the gain-of-function mutation *nhr-49(et7)*, which results in a single amino acid substitution near the ligand binding domain of NHR-49 and was shown to upregulate putative targets (Lee et al. 2016). We found that *nhr-49(et7)* animals exhibit increased survival to lethal heat stress, at a level comparable to that of *cbd-1(ok2913)* (Fig. 5A). Importantly, we did not observe any cumulative effect on heat stress survival in *nhr-49(et7);cbd-1(ok2913)* double mutants (Fig. 5A), indicating that these two mutations act within the same genetic pathway to promote stress resilience.

**Figure 5.**
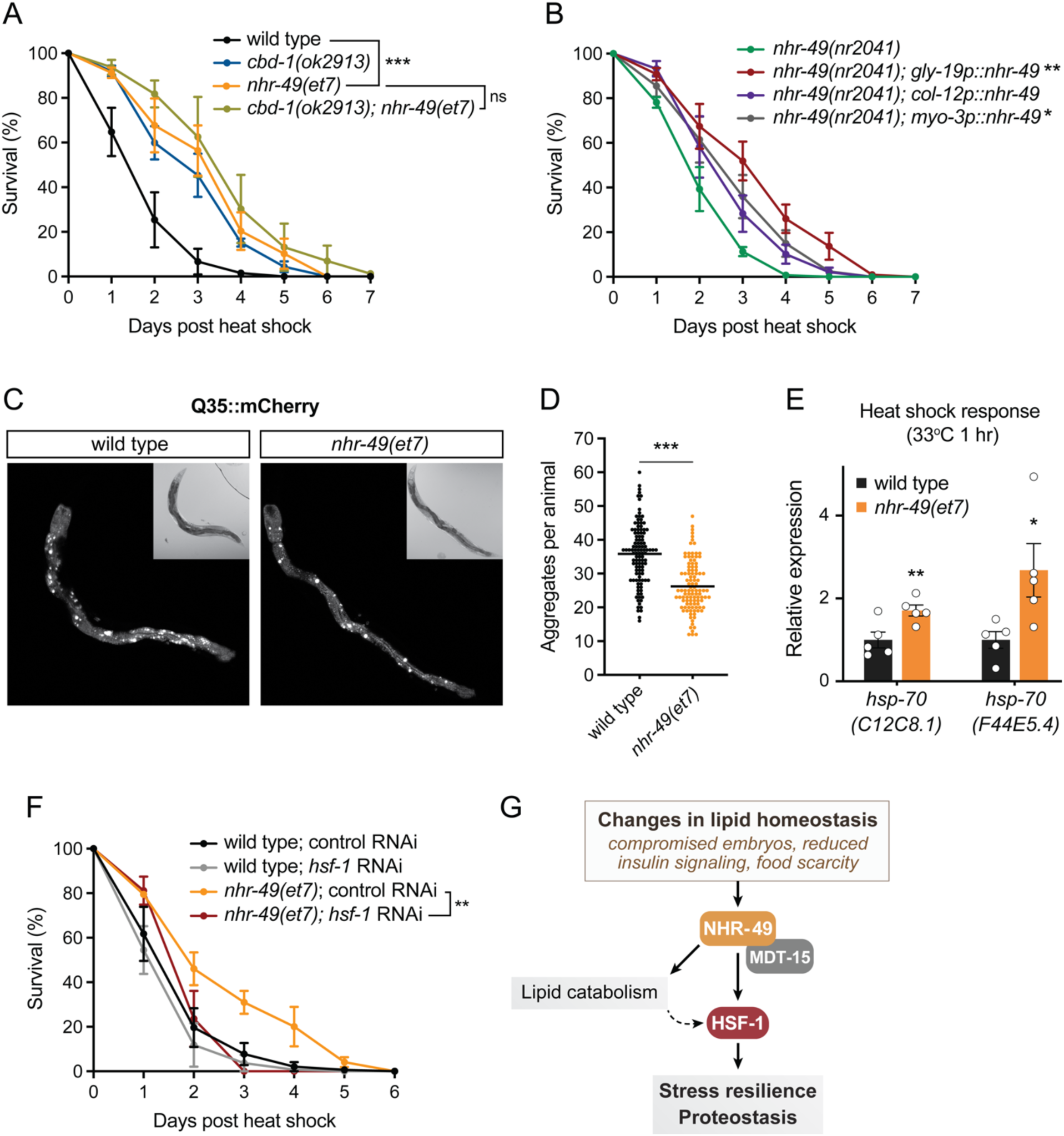
NHR-49 is sufficient to enhance stress resilience, proteostasis, and the HSR, and acts in an HSF-1-dependent manner. (A) Heat stress survival of wild type and *cbd-1(ok2913)* mutants crossed into the *nhr-49(et7)* gain-of-function allele (*n* = 3). (B) Heat stress survival of *nhr-49(nr2041)* and tissue-specific overexpression lines (*n* = 3). (C) Representative confocal images of day 5 adults expressing the Q35::mCherry construct in the intestine, with corresponding bright field images in inserts. (D) Quantification of Q35::mCherry aggregates in day 5 adults. Wild type, *n* = 112; *nhr-49(et7)*, *n* = 106. (E) Expression of heat-shock genes relative to *cdc-42* in wild-type animals exposed to 33°C heat shock for 1 hour (*n* = 5). (F) Heat stress survival of wild type or *nhr-49(et7)* animals grown on either control or *hsf-1* RNAi (*n* = 3). (G) Model for regulation of stress resilience and proteostasis by NHR-49 in response to changes in lipid homeostasis. Changes in lipid homeostasis, which can be caused by damage to the fertilized embryo, reduced insulin signaling, or food scarcity, activate NHR-49 which, together with its co-activator MDT-15, induces the expression of genes involved in lipid catabolism. NHR-49 activation also leads to potentiation of HSF-1 activity and enhancement of the HSR, resulting in improved stress resilience and proteostasis. Error bars represent SEM. Statistical significance based on two-way ANOVA (A, B, F) or unpaired *t*-test (D, E). (*) *P* < 0.05; (**) *P* < 0.01; (***) *P* < 0.001, (ns [nonsignificant]) *P* > 0.05.

We next asked whether overexpression of NHR-49 in individual somatic tissues was sufficient to impact stress resilience. For this, we used animals that overexpress the NHR-49::GFP protein from extrachromosomal arrays under the control of different tissue-specific promoters, namely *gly-19p* (intestine), *col-12p* (hypodermis), or *myo-3p* (muscle), in an otherwise *nhr-49(nr2041)* mutant background (Naim et al. 2021). We measured the ability of these strains to survive following a lethal heat stress and found that overexpression of NHR-49 in all three tissues tested improved survival compared to the *nhr-49(nr2041)* control, with the strongest effect being observed for intestinal-specific overexpression (Fig. 5B).

To determine whether NHR-49 activity also had beneficial effects in the face of chronic proteotoxic stress, we asked whether gain-of-function or intestinal overexpression affected the age-dependent aggregation of a polyglutamine model protein expressed in intestinal cells. For this, we used a strain that harbors an integrated array of a transgene consisting of a stretch of 35 glutamines fused to mCherry fluorescent protein (Q35::mCherry) and that is driven by an intestine-specific *vha-6* promoter. The animals exhibit mCherry fluorescence in intestinal cells which over time changes from diffuse distribution to visible foci that correspond to protein aggregates. Foci formation begins around day 3 of adulthood and gradually worsens as the animals get older (Fig. S3A, B). We found that the *nhr-49(et7)* gain-of-function mutation or *gly-19p::nhr-49* intestinal overexpression significantly decreased aggregate formation of Q35::mCherry, with a 30% reduction in foci numbers at day 5 of adulthood compared to the wild type control (Fig. 4C; Fig. S3C).

Therefore, increased NHR-49 activity is sufficient to improve organismal stress resilience and to limit age-dependent protein aggregation. This indicates that NHR-49 drives a transcriptional program that directly or indirectly promotes cellular resilience during both acute and chronic proteotoxic stress.

### NHR-49 enhances the HSR and acts via HSF-1 to improve stress resilience

We have previously shown that damage to the embryonic vitelline layer acts via HSF-1 to restore organismal stress resilience and the HSR in reproductive adults (Sala et al. 2020). HSF-1 is also known to be regulated downstream of insulin and GSC signaling (Hsu et al. 2003; Morley and Morimoto 2004; Shemesh et al. 2013; Labbadia and Morimoto 2015b). If HSF-1 activity is increased by NHR-49, we would expect higher levels of induction of the HSR. We tested this by monitoring transcripts corresponding to the canonical heat shock genes *hsp-70*(*C12C8.1)* and *hsp-70(F44E5.4),* both encoding members of the Hsp70 family, in *nhr-49(et7)* gain-of-function mutants following a mild heat stress of 33°C for one hour. We found that the induction of both Hsp70 encoding genes was higher in *nhr-49(et7)* compared to wild type animal by 1.7- and 2.7-fold, respectively (Fig. 5E). This effect was specific to heat stress conditions as we did not observe any effect of the *nhr-49(et7)* gain-of-function on the basal levels of *hsp70* mRNA, in contrast to putative NHR-49 targets *far-3* and *cyp-29A2* which were upregulated 4- and 10-fold, respectively (Fig. S4). This indicates that increased NHR-49 activity potentiates the ability of adult animals to induce the HSR in response to acute heat stress. We thus asked whether increased NHR-49 activity acts via the master regulator of the HSR, HSF-1, to promote stress resilience. We observed a complete suppression of the elevated heat stress survival of the *nhr-49(et7)* gain-of-function mutant when the animals were exposed to *hsf-1* RNAi (Fig. 5F), indicating that NHR-49 acts via HSF-1 to enhance stress resilience. Together, the data presented in this study suggests that NHR-49, through its lipid sensing and gene regulatory function, represents a link between lipid metabolism and regulation of the HSR via modulation of HSF-1 activity (Fig. 5G).

## Discussion

In this study, we show that NHR-49 facilitates the adaptation to conditions that modify lipid homeostasis by regulating the expression of genes involved in lipid metabolism, but also by rewiring the activity of the master regulator of the HSR, HSF-1, to enhance cellular resilience towards both acute and chronic proteotoxic stress (Fig. 5G). While the HSF-1-mediated HSR is an ancient pathway conserved throughout the eukaryotic kingdom, nuclear hormone receptors are only present in multicellular organisms. We posit that regulation of the HSR by NHR-49 serves to coordinate programs of survival as organismal strategies based on certain metabolic and reproductive cues that converge on lipid homeostasis.

The role of NHR-49 in lipid homeostasis is complex, as it is required to maintain lipid stores in adults, but also promotes FA breakdown via β-oxidation (Van Gilst et al. 2005a; Van Gilst et al. 2005b; Ratnappan et al. 2014). We find that NHR-49 is important for stress resilience in conditions that increase fat stores (*daf-2*, *cbd-1*, and *glp-1* mutants), but also conditions of food deprivation where lipid stores are decreased (Fig. 4). This indicates that stress resilience and the activity of NHR-49 are not directly influenced by the level of stored fat. This could be explained by the production of a similar metabolic signal in these conditions that may be a ligand for NHR-49. Another possibility is that the response to different lipid levels or content could be driven by interaction with different partners, as NHR-49 was shown to heterodimerize with NHR-66 and NHR-80 to drive distinct programs (Pathare et al. 2012).

NHR-49 was recently shown to also be responsive to changes in protein homeostasis during loss of HSF-1, affecting the expression of lipid metabolism genes and lifespan, but not thermotolerance (Watterson et al. 2022). This is consistent with our findings that NHR-49 loss-of-function has little to no impact on stress resilience in wild type animals, and that its activation in conditions that change lipid homeostasis promotes cellular resilience to proteotoxic stress. Since we show that NHR-49 is dependent on HSF-1 for this effect, it is then expected that during loss of HSF-1, the protective effects of NHR-49 on stress resilience were not observed (Watterson et al. 2022). Taken together, the findings of Watterson et al (Watterson et al. 2022) and this study establish NHR-49 as a two-way regulator connecting lipid homeostasis and proteostasis.

We find that the HSR is enhanced by NHR-49 gain-of-function specifically in stress conditions (Fig. 5, Fig. S4), suggesting that NHR-49 potentiates the intrinsic activity of HSF-1. The transcriptional activity of HSF1 is modulated by posttranslational modifications, as well as chromatin remodeling factors that favor binding at heat shock gene promoters (Holmberg et al. 2001; Guertin and Lis 2010; Gomez-Pastor et al. 2018; Takii et al. 2019). An important step in the activation of the HSR is the recruitment of Mediator by HSF1 at heat shock promoters (Park et al. 2001; Anandhakumar et al. 2016), which physically interacts with and recruits RNA polymerase II (Soutourina 2018). Direct interaction of NHR-49/MDT-15 with HSF-1 or binding to a nearby motif may thus facilitate the recruitment of the transcription machinery to heat shock genes during proteotoxic stress. Future efforts to map the binding sites of NHR-49 in the *C. elegans* genome will shed light on its mechanism of cooperation with HSF-1 during proteotoxic stress. NHR-49 may also promote the activity of HSF-1 indirectly. We show that the elevated stress resilience of the *cbd-1* vitelline mutant is hampered by *daf-22* knockdown (Fig. 2E), which suggests that changes in the peroxisomal β-oxidation pathway induced by NHR-49 may contribute to the elevated stress resilience phenotype.

Considering the conservation of these transcription factors, our finding that NHR-49 regulates HSF-1 and the HSR may be relevant to higher organisms. Interestingly, PPARα was shown to be required for the expression of a subset of heat stress-responsive genes in mouse liver, including genes previously shown to be HSF1-dependent (Vallanat et al. 2010). This suggests that the regulatory role of NHR-49/PPARα in the HSR may be conserved. Overall, our findings that the lipid sensing nuclear receptor NHR-49 acts as a critical regulator of cellular resilience to proteotoxic stress and proteostasis link two major cellular processes and form the basis for future studies to further understand the interconnections between lipid homeostasis and regulation of the HSR as organismal strategies to optimize fitness and survival.

## Materials and Methods

### *C. elegans* strains and maintenance

Standard *C. elegans* methods were used as previously described (Brenner 1974). Worms were maintained on solid Nematode Growth Medium (NGM) seeded with *E. coli* OP50 or for RNAi experiments, HT115 transformed with appropriate plasmids. All experiments were performed at 20 °C unless stated otherwise. Age synchronization was performed by timed egg laying in a one-to two-hour period for all experiments.

The following strains were used in this study: wild-type (N2 Bristol), VC2258 *cbd-1(ok2913) IV*, RT130 *pwIs23 [vit-2::GFP]*, STE68 *nhr-49(nr2041) I*, STE108 *nhr-49(et7) I*, AGP65 *nhr-49(nr2041) I;glmEx9 [pgly-19::nhr-49::GFP+pmyo-2 mCherry]*, AGP56 *nhr-49(nr2041) I; glmEx11 [Pcol-12::nhr-49::GFP + Pmyo-2::mCherry]*, AGP63 *nhr-49(nr2041) I;glmEx8 [pmyo-3::nhr-49::GFP + pmyo-2::mCherry]*. Strain VC2258 was generated by the *C. elegans* Gene Knockout Consortium and was backcrossed six times to N2 before use. STE108 was obtained from Stephan Taubert, and AGP65, AGP63, and AGP56 from Arjumand Ghazi. All other strains were either generated in our laboratory or obtained from the *Caenorhabditis* Genetics Center (CGC). RT130 was backcrossed three times to N2 before use to alleviate a slow growth phenotype.

The VC2258 strain was crossed to STE68 and STE108 to generate strains AM1238 and AM1237, respectively. Strain STE68 was also crossed to CB4037 and CB1370, resulting in strains AM1243 and AM1244.

A plasmid construct for the expression of Q35::mCherry was generated by inserting the sequence encoding Q35 upstream of *mCherry* in pGH8 (gift from Erik Jorgensen; Addgene plasmid #19359) and replacing the promoter with a 880-base pair sequence corresponding to the *vha-6* promoter using the NEBuilder HiFi DNA Assembly cloning kit. The resulting plasmid was injected into the gonad of adult N2 hermaphrodites at 10 ng/µl and fluorescent progeny was isolated to establish extrachromosomal lines. The array was integrated by UV irradiation and the resulting strain was backcrossed five times to N2 animals to generate strain AM1240 *rmIs406 [vha-6p::Q35::mCherry]*. The *rmIs406* allele was crossed with the *nhr-49(et7)* allele to generate strain AM1241, and with the *glmEx9* array to generate strain AM1242.

### RNA extraction and sequencing

Age-synchronized populations of at least 200 animals were grown at 20 °C until day 4 of adulthood and transferred to fresh plates every day to avoid contamination with progeny. Heat shock was performed on NGM plates wrapped with parafilm and submerged in a water bath at 33 °C for 1 hour. Non-treated and heat stressed samples were then collected into M9 media and snap frozen in liquid nitrogen. RNA was extracted using Trizol (Invitrogen) and purified with QIAGEN RNeasy MinElute RNA extraction kit with on-column DNase I digestion according to manufacturers’ instructions. Samples were prepared in biological triplicates.

Library preparation and sequencing were performed by Novogene (Shanghai, China). Briefly, 1 μg of RNA was used to generate sequencing libraries using NEBNext® UltraTM RNA Library Prep Kit for Illumina® (NEB, USA) following polyA selection. The library preparations were sequenced on an Illumina HiSeq 4000 instrument and 150 bp paired-end reads were generated.

To facilitate reproducible analysis, samples were processed using the publicly available nf-core/RNA-seq pipeline version 3.8.1 implemented in Nextflow 22.04.5.5708 using Singularity 3.8.1 with the minimal command nextflow run nf-core/rnaseq \ -r ‘3.8.1’ \ -profile nu_genomics -- genome ‘WBcel235’. Briefly, lane-level reads were trimmed using trimGalore! 0.6.7 and aligned to the WBcel235 genome described above using STAR 2.6.1d. Gene-level assignment was then performed using salmon 1.5.2.

### Differential expression analysis

Differential expression analysis (DEA) was performed using custom scripts in R version 4.1.1 using the DESeq2 version 1.34.0 framework. A “local” model of gene dispersion was employed as this better fit dispersion trends without obvious overfitting, and pairwise comparisons were performed by treating genotype and treatment (HS) as independent factors (~ genotype + treatment), as no significant interaction between these factors was observed for any gene. For pairwise comparisons, a combined factor of genotype and treatment was used (~ combined). Alpha was set at 0.05 for all DEA. Otherwise default settings were used. See code for details. High-level analysis was performed using custom scripts available in the nu-pulmonary/utils GitHub repository. GO term enrichment was then determined using Fisher’s exact test (classic mode) with FDR correction in topGO version 2.46.0, with org.Ce.eg.db version 3.14.0 as a reference. For KEGG enrichment analysis, an analogous approach was employed using manual Fisher’s exact tests with FDR correction against all annotated KEGG pathways in *C. elegans* using KEGGREST 1.34.0. Tissue and phenotype enrichment were determined using the Gene Set Enrichment Analysis tool available on Wormbase (Angeles-Albores et al. 2016). Transcriptomic data of *nhr-49(nr2041)* mutants from Watterson et al (Watterson et al. 2022) was obtained from the NCBI Gene Expression Omnibus (GEO) database (accession GSE199971) and analyzed using the same method and parameters.

### Heat stress survival assay

Synchronized animals (~50 per plate) were grown at 20 °C until either day 2 or day 4 of adulthood, transferred to new plates, sealed with parafilm, and incubated for 4 hours in a water bath at 35 °C. Worms were allowed to recover at 20 °C and survival was scored every 24 hours until all animals were dead. Animals were scored as dead in the absence of pharyngeal pumping or touch response, and animals with prolapsed gonads were censored from the analysis. Scoring was performed in a a blinded manner and the assay repeated at least three times per condition.

For stress survival assays following food deprivation, age-synchronized animals were grown to day 1 of adulthood (72 hours post-egg lay) and transferred to NGM plates without any bacterial food source. After 16 hours of food deprivation, the animals were transferred back onto OP50-seeded NGM plates and heat shocked for 4 hours as detailed above.

### Brood size

Young adults were singled onto OP50 seeded plates and transferred to fresh plates every 24 hours for 5 days. Following transfer, the plates containing the eggs were kept at 20 °C for 24 hours and viable progeny was counted.

### Oil red O staining and quantification

Oil red O staining of neutral lipids was performed in fixed animals as previously described (Escorcia et al. 2018). Briefly, age-synchronized day 2 animals were washed in PBS supplemented with 0.01% Triton X-100 (PBST) and incubated in 40% isopropanol for 3 minutes. Oil red O (Sigma O0625) resuspended in 60% isopropanol was added to each tube and worms were rocked at room temperature for 2 hours. After a 30-minute wash in PBST, animals were mounted on slides and imaged on an Olympus BX53 microscope using a Micropublisher color camera. Color images were quantified in ImageJ as previously described (O’Rourke et al. 2009) by measuring within each worm object the level of intensity in the red channel in areas with excess red intensity compared to blue and green signals, and normalizing to the area of the object.

### Triglyceride quantification

Approximately 200 animals were collected on day 2 of adulthood in water and washed extensively. The worms were resuspended in lysis buffer (25 mM Tris HCl (pH 7.4), 1 mM EDTA, 1% Triton X-100) and sonicated with a Bioruptor 30 seconds on, 30 seconds off at maximum power for 10 minutes. The lysate was cleared of debris with a brief spin and protein concentration was estimated using the Bradford assay. Lipids were dissolved by adding 5% NP40 to the samples and slowly heating them to 100°C, followed by cooling at room temperature. This was preformed twice before the lysate was cleared by centrifugation. The supernatant was then used for the Triglyceride Quantification Fluorometric assay (Sigma MAK266) per the manufacturer’s instructions. Experiments were performed in biological triplicates for each condition examined.

### Fatty acid composition analysis

Fatty acid composition was determined through direct transesterification of worm extracts to generate fatty acid methyl ester (FAMEs). Approximately 1000 age-synchronized worms were added to 1ml of 2.5% sulfuric acid in methanol and heated for one hour at 70°C. FAMES were extracted with hexane and injected into a gas chromatography/mass spectrometry (GC/MS) equipped with a SP-2380 column for separation and analysis (Harrison and Watts 2022).

### RNAi treatments

For RNAi-mediated knockdown of indicated genes, indicated strains were synchronized by egg laying on *E. coli* strain HT115(DE3) containing the appropriate RNAi vectors obtained from the Ahringer RNAi library, with L4440 as the empty vector control (Kamath et al. 2003). RNAi bacteria were grown overnight, and cultures were induced with 5 mM IPTG for 3 hours.

### Real-time qPCR

RNA extraction was performed on populations of at least 100 animals per replicate, using TRIzol (Invitrogen) and RNA purified with QIAGEN RNeasy MinElute columns as described previously (Labbadia and Morimoto 2015b). mRNA was reverse transcribed using the iScript cDNA Synthesis Kit (BioRad) and real-time quantitative PCR was performed using iQ SYBR Green Supermix (BioRad) in a BioRad CFX384 Real-Time PCR system. Relative expression was determined from cycle threshold values using the standard curve method and the expression of genes of interest was normalized to *rpb-2*. The primers used are listed in Table S1.

### Fluorescence microscopy

Age-synchronized animals were mounted on 3% agarose pads in 2 mM levamisole in M9 buffer and imaged with a Zeiss LSM 800 confocal microscope using a ×10 or ×20 objective lens and the Zen imaging software, and maximum intensity projection images were generated from Z stacks using ImageJ software.

### Quantification of polyglutamine aggregates

Populations of 30 age-synchronized adult animals were transferred to fresh plates every day, and the number of Q35::mCherry aggregates was visually determined in live animals by counting large, bright fluorescent foci using a fluorescence stereomicroscope. Experiments were repeated three times.

## Data availability

Raw transcriptomic data files generated for this study have been deposited on the SRA database (http://www.ncbi.nlm.nih.gov/sra) under the accession PRJNA948034.

## Acknowledgments

We thank Stefan Taubert, Arjumand Ghazi, and Erik Jorgensen for sharing strains and plasmids, and Jian Li for critical reading and comments on the manuscript. We also thank the Keck Biophysics, High Throughput Analysis, and Biological Imaging facilities at Northwestern University for access to equipment. This work was supported by grants from the NIH (RF1AG057296, P01AG054407) and the Daniel F. and Ada L. Rice Foundation to R.I.M., NIH grant T32AG020506-18 to R.A.G., NIH grant R01GM13883 to J.L.W, and a postdoctoral fellowship from the American Federation for Aging Research to L. C. B..

## Author contributions

AJS conceived the study and designed the experiments with input from RIM; AJS, GI, RMB, CM, and JW performed the experiments; LCB contributed reagents; AJS, RAG, and JW analyzed the data; AJS wrote the manuscript with input from RIM and LCB.

## References

Amrit FR, Steenkiste EM, Ratnappan R, Chen SW, McClendon TB, Kostka D, Yanowitz J, Olsen CP, Ghazi A. 2016. DAF-16 and TCER-1 Facilitate Adaptation to Germline Loss by Restoring Lipid Homeostasis and Repressing Reproductive Physiology in C. elegans. PLoS Genet 12: e1005788.

Anandhakumar J, Moustafa YW, Chowdhary S, Kainth AS, Gross DS. 2016. Evidence for Multiple Mediator Complexes in Yeast Independently Recruited by Activated Heat Shock Factor. Mol Cell Biol 36: 1943–1960.

Angeles-Albores D, RY NL, Chan J, Sternberg PW. 2016. Tissue enrichment analysis for C. elegans genomics. BMC Bioinformatics 17: 366.

Antebi A. 2013. Regulation of longevity by the reproductive system. Exp Gerontol 48: 596–602.

Balch WE, Morimoto RI, Dillin A, Kelly JW. 2008. Adapting proteostasis for disease intervention. Science 319: 916–919.

Brenner S. 1974. The genetics of Caenorhabditis elegans. Genetics 77: 71–94.

Doering KRS, Cheng X, Milburn L, Ratnappan R, Ghazi A, Miller DL, Taubert S. 2022. Nuclear hormone receptor NHR-49 acts in parallel with HIF-1 to promote hypoxia adaptation in Caenorhabditis elegans. Elife 11.

Escorcia W, Ruter DL, Nhan J, Curran SP. 2018. Quantification of Lipid Abundance and Evaluation of Lipid Distribution in Caenorhabditis elegans by Nile Red and Oil Red O Staining. J Vis Exp.

Flatt T, Min KJ, D’Alterio C, Villa-Cuesta E, Cumbers J, Lehmann R, Jones DL, Tatar M. 2008. Drosophila germ-line modulation of insulin signaling and lifespan. Proc Natl Acad Sci U S A 105: 6368–6373.

Fontana L, Partridge L. 2015. Promoting health and longevity through diet: from model organisms to humans. Cell 161: 106–118.

Goh GYS, Winter JJ, Bhanshali F, Doering KRS, Lai R, Lee K, Veal EA, Taubert S. 2018. NHR-49/HNF4 integrates regulation of fatty acid metabolism with a protective transcriptional response to oxidative stress and fasting. Aging Cell 17: e12743.

Gomez-Pastor R, Burchfiel ET, Thiele DJ. 2018. Regulation of heat shock transcription factors and their roles in physiology and disease. Nat Rev Mol Cell Biol 19: 4–19.

Gonzalez DP, Lamb HV, Partida D, Wilson ZT, Harrison MC, Prieto JA, Moresco JJ, Diedrich JK, Yates JR, 3rd, Olson SK. 2018. CBD-1 organizes two independent complexes required for eggshell vitelline layer formation and egg activation in C. elegans. Dev Biol 442: 288–300.

Guertin MJ, Lis JT. 2010. Chromatin landscape dictates HSF binding to target DNA elements. PLoS Genet 6: e1001114.

Hansen M, Flatt T, Aguilaniu H. 2013. Reproduction, fat metabolism, and life span: what is the connection? Cell Metab 17: 10–19.

Harrison HH, Watts JL. 2022. Lipid Extraction and Analysis. Methods Mol Biol 2468: 271–281.

Hipp MS, Kasturi P, Hartl FU. 2019. The proteostasis network and its decline in ageing. Nat Rev Mol Cell Biol 20: 421–435.

Holmberg CI, Hietakangas V, Mikhailov A, Rantanen JO, Kallio M, Meinander A, Hellman J, Morrice N, MacKintosh C, Morimoto RI et al. 2001. Phosphorylation of serine 230 promotes inducible transcriptional activity of heat shock factor 1. EMBO J 20: 3800–3810.

Hsin H, Kenyon C. 1999. Signals from the reproductive system regulate the lifespan of C. elegans. Nature 399: 362–366.

Hsu AL, Murphy CT, Kenyon C. 2003. Regulation of aging and age-related disease by DAF-16 and heat-shock factor. Science 300: 1142–1145.

Johnston WL, Krizus A, Dennis JW. 2010. Eggshell chitin and chitin-interacting proteins prevent polyspermy in C. elegans. Curr Biol 20: 1932–1937.

Kamath RS, Fraser AG, Dong Y, Poulin G, Durbin R, Gotta M, Kanapin A, Le Bot N, Moreno S, Sohrmann M et al. 2003. Systematic functional analysis of the Caenorhabditis elegans genome using RNAi. Nature 421: 231–237.

Kenyon C, Chang J, Gensch E, Rudner A, Tabtiang R. 1993. A C. elegans mutant that lives twice as long as wild type. Nature 366: 461–464.

Labbadia J, Morimoto RI. 2015a. The biology of proteostasis in aging and disease. Annu Rev Biochem 84: 435–464.

Labbadia J, Morimoto RI. 2015b. Repression of the Heat Shock Response Is a Programmed Event at the Onset of Reproduction. Mol Cell 59: 639–650.

Lee K, Goh GY, Wong MA, Klassen TL, Taubert S. 2016. Gain-of-Function Alleles in Caenorhabditis elegans Nuclear Hormone Receptor nhr-49 Are Functionally Distinct. PLoS One 11: e0162708.

Leone TC, Weinheimer CJ, Kelly DP. 1999. A critical role for the peroxisome proliferator-activated receptor alpha (PPARalpha) in the cellular fasting response: the PPARalpha-null mouse as a model of fatty acid oxidation disorders. Proc Natl Acad Sci U S A 96: 7473–7478.

Li J, Labbadia J, Morimoto RI. 2017. Rethinking HSF1 in Stress, Development, and Organismal Health. Trends Cell Biol 27: 895–905.

Morley JF, Morimoto RI. 2004. Regulation of longevity in Caenorhabditis elegans by heat shock factor and molecular chaperones. Mol Biol Cell 15: 657–664.

Naim N, Amrit FRG, Ratnappan R, DelBuono N, Loose JA, Ghazi A. 2021. Cell nonautonomous roles of NHR-49 in promoting longevity and innate immunity. Aging Cell 20: e13413.

O’Rourke EJ, Soukas AA, Carr CE, Ruvkun G. 2009. C. elegans major fats are stored in vesicles distinct from lysosome-related organelles. Cell Metab 10: 430–435.

Park JM, Werner J, Kim JM, Lis JT, Kim YJ. 2001. Mediator, not holoenzyme, is directly recruited to the heat shock promoter by HSF upon heat shock. Mol Cell 8: 9–19.

Pathare PP, Lin A, Bornfeldt KE, Taubert S, Van Gilst MR. 2012. Coordinate regulation of lipid metabolism by novel nuclear receptor partnerships. PLoS Genet 8: e1002645.

Ratnappan R, Amrit FR, Chen SW, Gill H, Holden K, Ward J, Yamamoto KR, Olsen CP, Ghazi A. 2014. Germline signals deploy NHR-49 to modulate fatty-acid beta-oxidation and desaturation in somatic tissues of C. elegans. PLoS Genet 10: e1004829.

Sala AJ, Bott LC, Brielmann RM, Morimoto RI. 2020. Embryo integrity regulates maternal proteostasis and stress resilience. Genes Dev 34: 678–687.

Sala AJ, Bott LC, Morimoto RI. 2017. Shaping proteostasis at the cellular, tissue, and organismal level. J Cell Biol 216: 1231–1241.

Sala AJ, Morimoto RI. 2021. Protecting the future: balancing proteostasis for reproduction. Trends Cell Biol.

Shemesh N, Shai N, Ben-Zvi A. 2013. Germline stem cell arrest inhibits the collapse of somatic proteostasis early in Caenorhabditis elegans adulthood. Aging Cell 12: 814–822.

Shi X, Li J, Zou X, Greggain J, Rodkaer SV, Faergeman NJ, Liang B, Watts JL. 2013. Regulation of lipid droplet size and phospholipid composition by stearoyl-CoA desaturase. J Lipid Res 54: 2504–2514.

Shmookler Reis RJ, Xu L, Lee H, Chae M, Thaden JJ, Bharill P, Tazearslan C, Siegel E, Alla R, Zimniak P et al. 2011. Modulation of lipid biosynthesis contributes to stress resistance and longevity of C. elegans mutants. Aging (Albany NY) 3: 125–147.

Soutourina J. 2018. Transcription regulation by the Mediator complex. Nat Rev Mol Cell Biol 19: 262–274.

Takii R, Fujimoto M, Matsumoto M, Srivastava P, Katiyar A, Nakayama KI, Nakai A. 2019. The pericentromeric protein shugoshin 2 cooperates with HSF1 in heat shock response and RNA Pol II recruitment. EMBO J 38: e102566.

Tatar M, Bartke A, Antebi A. 2003. The endocrine regulation of aging by insulin-like signals. Science 299: 1346–1351.

Taubert S, Van Gilst MR, Hansen M, Yamamoto KR. 2006. A Mediator subunit, MDT-15, integrates regulation of fatty acid metabolism by NHR-49-dependent and -independent pathways in C. elegans. Genes Dev 20: 1137–1149.

Vallanat B, Anderson SP, Brown-Borg HM, Ren H, Kersten S, Jonnalagadda S, Srinivasan R, Corton JC. 2010. Analysis of the heat shock response in mouse liver reveals transcriptional dependence on the nuclear receptor peroxisome proliferator-activated receptor alpha (PPARalpha). BMC Genomics 11: 16.

Van Gilst MR, Hadjivassiliou H, Jolly A, Yamamoto KR. 2005a. Nuclear hormone receptor NHR-49 controls fat consumption and fatty acid composition in C. elegans. PLoS Biol 3: e53.

Van Gilst MR, Hadjivassiliou H, Yamamoto KR. 2005b. A Caenorhabditis elegans nutrient response system partially dependent on nuclear receptor NHR-49. Proc Natl Acad Sci U S A 102: 13496–13501.

Wang MC, O’Rourke EJ, Ruvkun G. 2008. Fat metabolism links germline stem cells and longevity in C. elegans. Science 322: 957–960.

Wani KA, Goswamy D, Taubert S, Ratnappan R, Ghazi A, Irazoqui JE. 2021. NHR-49/PPAR-alpha and HLH-30/TFEB cooperate for C. elegans host defense via a flavin-containing monooxygenase. Elife 10.

Watterson A, Arneaud SLB, Wajahat N, Wall JM, Tatge L, Beheshti ST, Mihelakis M, Cheatwood NY, McClendon J, Ghorashi A et al. 2022. Loss of heat shock factor initiates intracellular lipid surveillance by actin destabilization. Cell Rep 41: 111493.

Wilson DM, 3rd, Cookson MR, Van Den Bosch L, Zetterberg H, Holtzman DM, Dewachter I. 2023. Hallmarks of neurodegenerative diseases. Cell 186: 693–714.

