## Supplemental Figures and Table for "Nuclear receptor signaling via NHR-49/MDT-15 regulates stress resilience and proteostasis in response to reproductive and metabolic cues"

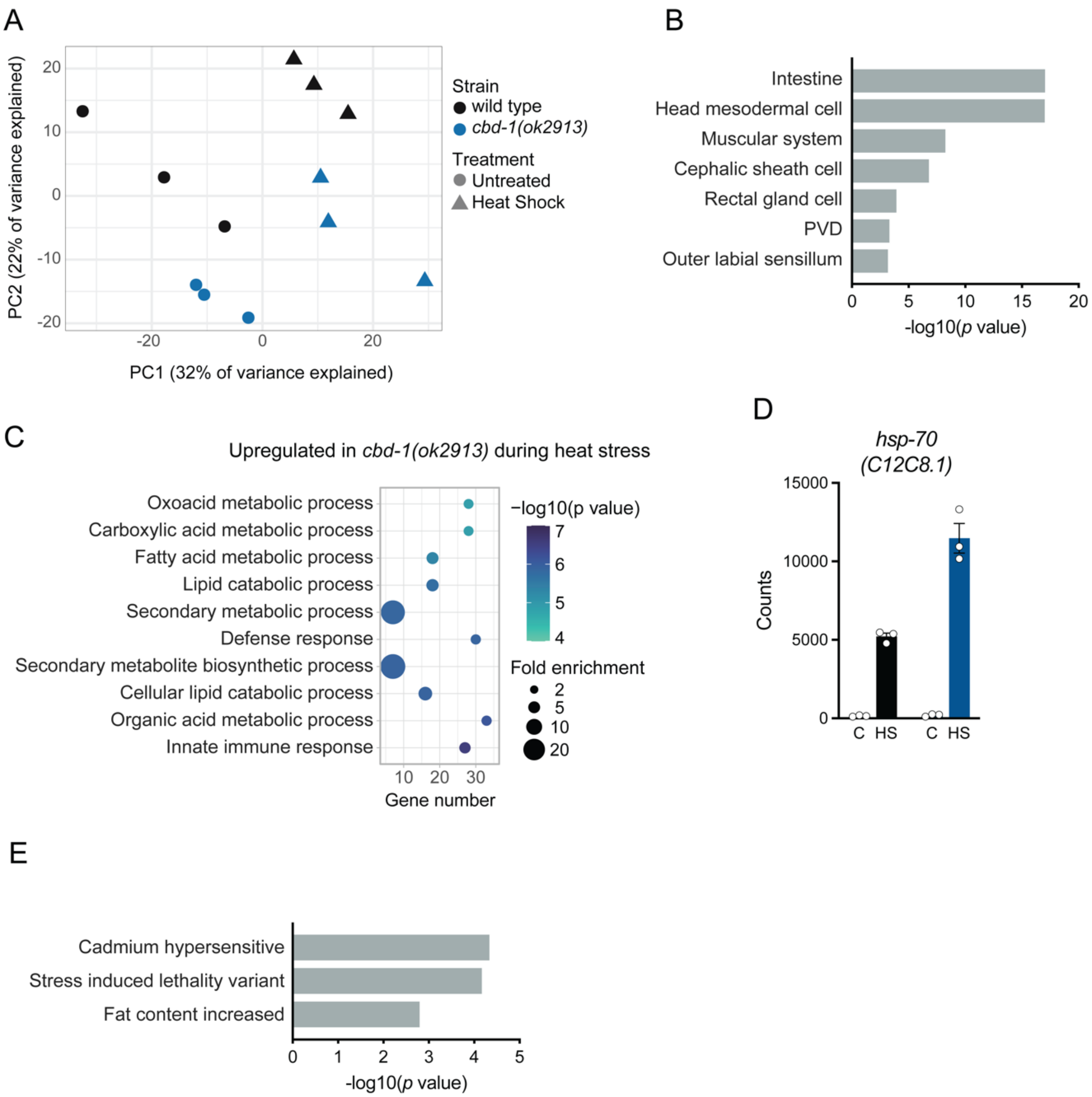


**Figure S1. Gene expression analysis of *cbd-1(ok2913)***

(A) Principal component analysis of RNA sequencing data. (B) Tissue enrichment analysis of genes differentially expressed in *cbd-1(ok2913)* mutants. (C) Gene Ontology enrichment analysis in genes upregulated in *cbd-1(ok2913)* compared to wild type following a 1-hour heat stress at 33ºC. (D) Expression of *hsp-70* (*C12C8.1*) in wild type (black) and *cbd-1(ok2913)* animals in control conditions (C) and following heat stress (HS). (E) Phenotype enrichment analysis of genes differentially expressed in *cbd-1(ok2913) vs* wild type animals.

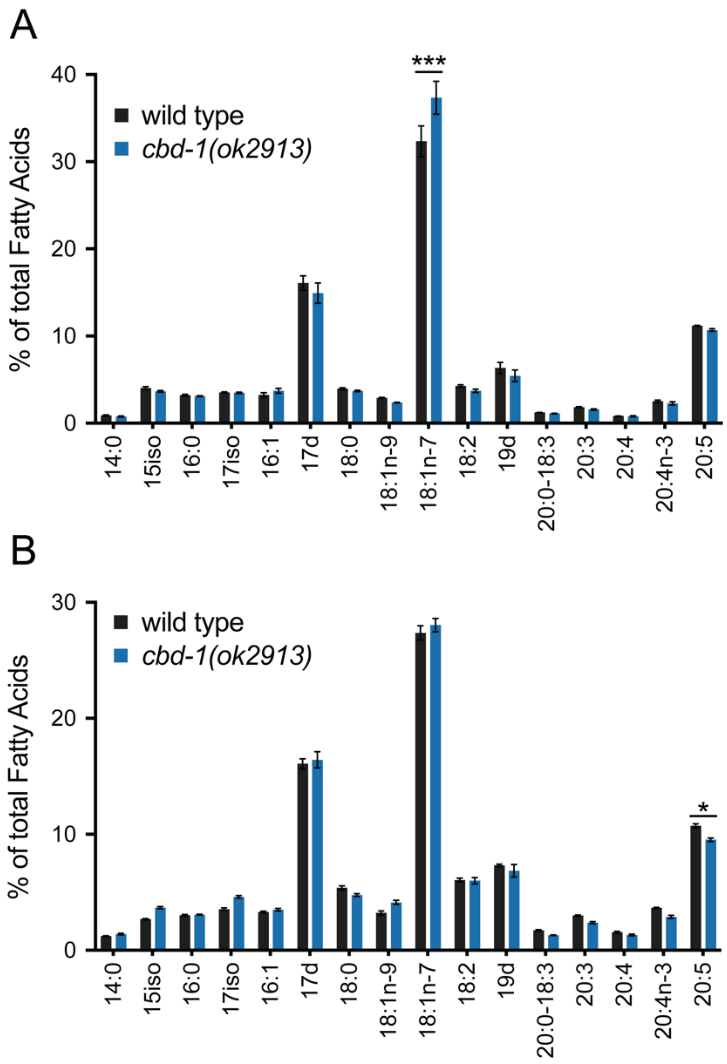


**Figure S2. Profiling of relative FA content in wild type and *cbd-1(ok2913)* animals**

Relative fatty acid composition of gravid day 1 (A) and day 4 (B) animals ($n$ = 3). Error bars represent SEM. Statistical significance based on two-way ANOVA followed by Sidak correction. (*) $P < 0.05$; (***) $P < 0.001$.

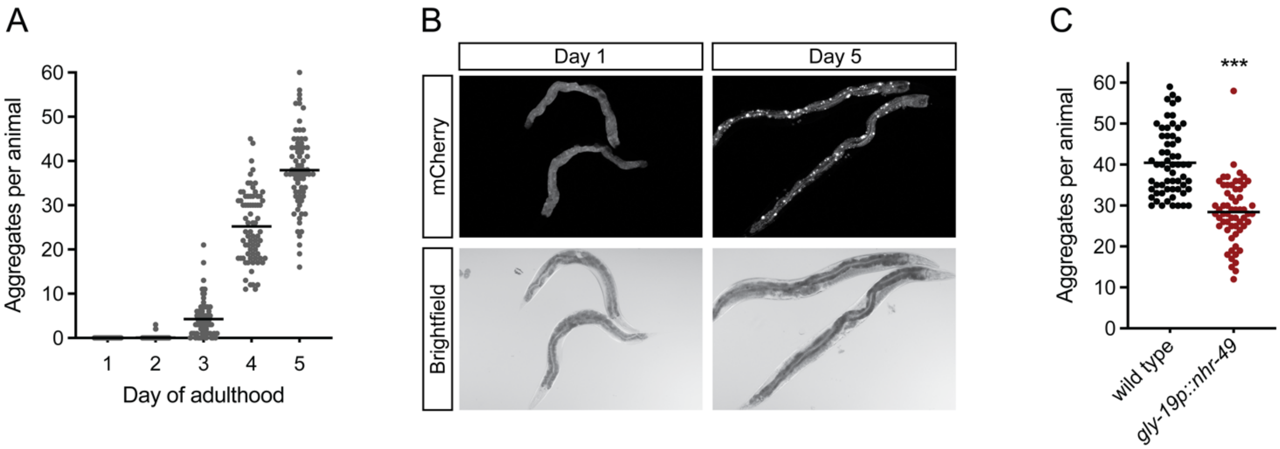


**Figure S3. Characterization of Q35::mCherry localization in intestinal cells**

(A) Quantification of Q35::mCherry aggregates on the indicated days of adulthood. Day 1, $n$ = 83; day 2, $n$ = 83; day 3, $n$ = 80; day 4, $n$ = 74; day 5, $n$ = 72. (B) Representative confocal images of day 1 and day 5 animals showing the relocalization of Q35::mCherry from a diffuse distribution to foci over time. (C) Quantification of Q35::mCherry aggregates in day 5 adults ($n$ = 60). Statistical significance based on unpaired $t$-test. (***) $P < 0.001$.

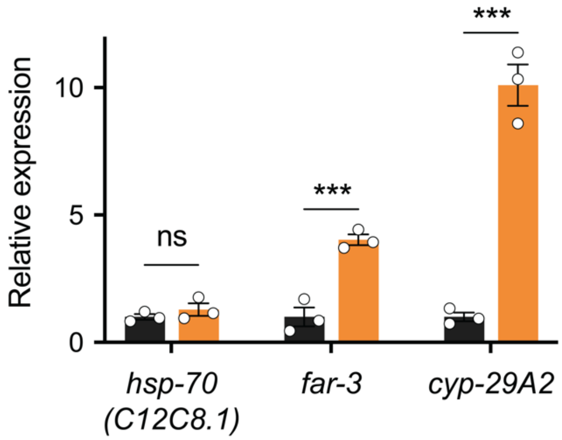


**Figure S4. Gene expression in the gain of function *nhr-49(et7)* mutant**

Expression of *C12C8.1*, *far-3*, and *cyp-29A2* relative to *rpb-2* in wild type (black) and *nhr-49(et7)* (orange) animals, normalized to wild type ($n$ = 3).

| **Target** | **Forward (5'-3')** | **Reverse (5'-3')** |
|---|---|---|
| *rpb-2* | AACTGGTATTGTGGATCAGGTG | TTTGACCGTGTCGAGATGC |
| *far-3* | CAAGTTGCTGCATACAAGGCA | CTTGGCGACTCCTCCGAAAT |
| *cyp-29A2* | GGGCAACTGGGTATAAAGCTCA | TCTCCGGAATCATGAGCAGC |
| *cdc-42* | GGTTGCTCCAGCTTCATTC | AACAAGAATGGGGTCTTTGA |
| *C12C8.1* | CTACATGCAAAGCGATTGGA | GGCGTAGTCTTGTTCCCTTC |
| *F44E5.4* | TGATACCCATCTCGGAGGAG | GTGGATTGGGTGAAATGTCC |

**Supplemental Table S1.** List of qPCR primers
